# ZnO nanoparticles and SWCNT induced general stress response pathway in HepG2 cells at non-cytotoxic doses revealed by RNA sequencing

**DOI:** 10.1101/2022.09.16.508235

**Authors:** Deepti Mittal, Syed Azmal Ali, Gautam Kaul

**Affiliations:** Nanoparticle studies lab, Animal Biochemistry Division, National Dairy Research Institute, Haryana, India; Cell Biology and Proteomics Lab, Animal Biotechnology Center, National Dairy Research Institute, Haryana, India

**Keywords:** Zinc Oxide, Nanoparticles, Single-Walled Carbon Nanotubes, Transcriptomics, Toxicity, Reactive oxygen species

## Abstract

Nanoparticles (NPs) are important in a variety of sectors, including disease diagnostics, medicine, nutrition, and many other industries. The risk of human exposure demands an early evaluation of both the basic dynamics of NPs’ interaction with biological systems and their potential consequences. Deciphering these occurrences will provide critical information regarding the health hazards and safety advantages associated with next-generation nanoformulations in clinical practice. We examined the HepG2 cell line in a systematic manner to determine the cellular response to single-walled carbon nanotubes (SWCNTs) and zinc oxide (ZnO) NPs. With the use of high-throughput transcriptomic methods, we found that both NPs induce comparable dysregulation of the endocytic and proteasomal complex genes in liver hepatocellular carcinoma cells, at levels (> 80 percent cell viability) that do not cause over-toxicity at early incubation period (6 h). SWCNT and ZnO NPs were shown to enter cells through clathrin-mediated pathways, affecting cytoskeleton gene expression, DNA damage and repair, protein ubiquitination, and cell transcriptional machinery. Our findings indicate that early response strategies activate stress-related mechanisms. Finally, this method for studying nanomaterial–cell interactions demonstrates how changes in the transcriptome profile may predict downstream consequences even at doses that do not cause acute toxicity.

## 1. Introduction

The recent world urbanisation leads to the excess use of nanotechnology, albeit of their tremendous implication at the industrial level; still, the mechanism of their action is poorly understood. Nanoparticles (NPs) are distinguished by their diverse physical and chemical characteristics, which include a wide range of sizes, morphologies, surface area, and surface activity. Two important classes of NPs: Metal oxide (ZnO) and carbon-based high aspect ratio materials (SWCNT), are produced with the highest global levels and utilised in a wide variety of applications. The presence of mystical structural properties in multi-walled carbon nanotubes (MWCNT) and single-walled carbon nanotubes (SWCNT) make them novel and attractive candidates to be used as chemical sensors (Meyyapan, 2016; Ali et al., 2021), superconductors (Futaba et al., 2006), a delivery vehicle for drugs, biomedical (Rafeeqi and Kaul, 2010) and agricultural (Patel et al., 2020) applications. Similarly, ZnO NPs have been utilised in various consumer products such as sunscreens for their excellent UV absorbing capacity (Mohammed et al., 2019), food packaging due to their great antimicrobial activity (Siddiqi et al., 2018) and as a potent antiviral agent (Mittal and Ali, 2022). It has also been realised to have growth-promoting and disease-preventing effects on various plants such as tomatoes and eggplants (Mittal et al., 2020; Elmer et al., 2016). The ever increasing commercial application and extensive utilisation of engineered NPs prompted the scientific community to identify potential impacts, including their toxicological property on human health and the environment.

The continuously evolving literature using *in vitro* and *in vivo* testing models revealed that NPs are responsible for generating reactive oxygen species (ROS), toxic ions shredding, lipid peroxidation, and genotoxicity. For instance, ZnO NPs are observed to be harmful to both *in vitro* (MRC-5 cell line) and *in vivo* (Drosophila) models, resulting in a significant increase in oxidative stress, decreased cellular viability genotoxicity and impairment of egg-to-adult viability (Ng et al., 2017). Also, SWCNT treated MRC-5 cells has shown a reduction in cell viability, significantly increased apoptosis and necrosis (Ong et al., 2017). In this context, the primary question arises whether this toxicological behaviour is based on the type of NPs or because of common characteristics shared by diverse NPs. On this confusion, the previously provided information regarding the safety aspects of these complex NPs are not acceptable and utilising the already established protocols may give misleading results. Furthermore, the recent report showed that the conventional assays used for the toxicity testing of drugs and soluble chemicals are not perfect for NPs because they tend to interfere with the components of these assays (Guadagini et al., 2015).

Furthermore, cytotoxicity analysis by conventional techniques does not provide the overall effect induced by NPs at the nano-bio interface. However, the detailed animal-based studies could not be applied to many NPs for their safety assessment. Instead, they are costly, labour intensive, and cannot be used as the primary screening platform. Therefore, new and highly robust testing strategies are urgently required to provide innumerable interactions of NPs with the biological system (Nel et al., 2013). Thus a great deal of effort must be put forth to identify the specific biomarker using high-throughput “omics” based technology. It also helps to uncover the potential mechanism induced by the NPs stress in the biological system. In this way, transcriptomic profiling is a powerful tool that can be utilised to evaluate the biological interaction for gene expression perturbations (Fröhlich et al., 2017). Furthermore, the previously used “omics” based microarray technique is confined to a limited pre-prescribed set of genes, unlike next-generation sequencing (NGS) that can directly sequence millions of transcripts without any pre-conceived idea of the genes and genome. Hence, it provides new avenues towards probing many pathways and molecular responses altogether (Carrow et al., 2018; Sun et al., 2017; Lucafò et al., 2013). Recently our lab has performed proteomics analysis that demonstrated dysregulation of proteins related to cytoskeleton alteration and thus affects cellular morphology by the effect of ZnO NPs, MWCNT and MSN (mesoporous silica nanoparticles) (Yadav et al., 2021). Thus this study provides enough support to move further with the omics analysis as a basis to uncover the cell-NPs interaction at a molecular level. Additionally our another study was based on the transcriptomic analysis for the effect of MCN (Mesoporous carbon nanoparticles) and MSN in liver hepatoma cells that describes the regulation of common molecular pathways during initial interaction with the cellular system. These NPs perturbed pathways including DNA damage, repair, cell cycle regulation, transcription and ubiquitination are common among both MCN and MSN (Mittal et al., 2022).

Previously, safety assessment studies that have been focussed using NGS mostly rely on the high dose exposure conditions that do not depict the realistic scenario. Recent efforts has been put forward to start assessing the toxicological effect of NPs considering the doses at which humans could be exposed in real life conditions (Kampfer et al., 2021). Understanding the toxicological effects of each NP type is critical for any prediction of their immediate and long-term risks for humans and ecosystems. Therefore, our study aimed to identify the low-dose and non-cytotoxic effects of ZnO NPs and SWCNT to get deeper insights into their cellular interaction.

## 2. Materials and Methods

### 2.1. Chemicals and materials

ZnO NPs dispersion (Cas. No. 1314-13-2; purity) and SWCNT powder (Cat. No. 704121-250MG), penicillin-streptomycin solution (Cat. No. P4333), MTT dye (Cat. No. M5655-1G), Neutral red dye (Cat. No. N4638-1G) and Trypan Blue dye (Cat. No. T6146-100G) were purchased from Sigma Aldrich (St. Louise, MO). Fetal bovine serum (FBS, Cat. No. CCS-500-SA-U), Dulbecco’s Modified Eagle’s Medium (DMEM) 1X with 4.5 g/L glucose, 4.0 mM L-glutamine, and sodium pyruvate (Cat. No. CC3004.05L), Dimethylsulphoxide (DMSO, Cat. No. PG-1320-500mL) all were purchased from CellClone™Genetix Biotech Asia Pvt Ltd. India. Dulbecco’s phosphate buffer saline (DPBS, Cat. No.SH30013.02) without calcium and magnesium was obtained from Hyclone Laboratories (Logan, UT).

### 2.2. Cell culture and exposure of nanoparticles

HepG2 cells were procured from National Centre for Cell Sciences (NCCS), Pune, India, and further grown in DMEM 1X medium supplemented with 10% FBS, 1% streptomycin and penicillin solution, in 25 cm^2^ flask, at 37**°**C in 5% CO_2_ humified atmosphere. The cells were used between 15-20 passages for each experiment. HepG2 cells were plated at a density of 3×10^5^ cells/ml in a 96 well plate and allowed to attach for 24 h before the treatment of NPs. The NPs were suspended in PBS to make the stock solution of 5 mg/ml and diluted to desired concentrations. The stock solution of NPs was sonicated on an ice bath for 10 min before giving treatment to the cells using a Branson ultrasonicator with a 13 mm probe diameter.

### 2.3. MTT assay

The cell viability was assessed by utilising the MTT assay according to Mosmann (1983) method with some modifications (Mosmann, 1983). First, the HepG2 cells were exposed to the NPs at the indicated concentrations for the different incubation periods. The medium was then removed and replaced with 10 µl of MTT [3-(4, 5-dimethylthiazoyl-2-yl)-2,5-diphenyltetrazolium bromide] dye (5 mg/ml) prepared in PBS and incubated for 4 h at 37**°**C. After incubation, the resulting formazan crystals were dissolved by adding 100 µl of dimethylsulphoxide, and the absorbance was measured at 570 nm in a multiplate reader. Three independent experiments were performed.

### 2.4. Neutral Red Uptake assay

The neutral red uptake (NRU) assay was performed according to Repetto et al with slight modification (Repetto et al., 2008). After the exposure of NPs, the medium was discarded, and 100 µl of neutral red dye (40 µg/ml) dissolved in serum-free medium was added to each well in a 96 well plate. After the desired incubation time was over, the excess dye solution was removed, and 150 µl of destaining solution (50% ethanol, 49% deionised water and 1% glacial acetic acid) was added. The plates were gently shaken, and absorbance was measured at 540 nm on an Infinite M200 multi-well plate reader (Tecan, Durham, USA). Three independent experiments were performed.

### 2.5. Trypan blue dye exclusion method

Trypan blue assay relies on the accumulation of dye in dead cells but is excluded from live cells and is the most commonly used procedure for cell viability assessment (Ishiyama et al., 1996; Strober, 2015). For trypan blue assay, 2 × 10^5^ cells were seeded in 24 well plate in 2 mL of complete culture medium and allowed to adhere for 24 h. The cells were then treated with ZnO NPs and SWCNT in a dose and time-dependent manner. After the exposure time, the cells were thoroughly washed with PBS, followed by incubation with trypsin and pelleted. These pelleted cells were dispersed in a fresh medium along with an equal volume of trypan blue (10 µl). Finally, the viable cells were counted using a Neubauer haemocytometer, where cell viability was assessed by counting both live and dead cells. The cell viability was calculated based on a triplicate of experiments.

### 2.6. Reactive oxygen species measurement with DCFH-DA assay

Intracellular ROS level was measured using dichlorodihydrofluorescein diacetate (DCFH-DA) assay (Wang and Joseph, 1999). Briefly, the HepG2 cells were seeded in 96 well black plates and incubated with ZnO NPs and SWCNT (0, 1, 2, 5, 10, 20, 25 and 50 µg/ml) for 1, 3, 6 and 12 h. After exposure, the cells were washed with PBS and loaded with 10 µM DCFH-DA dye in PBS for 40 min at 37°C. After that, the cells were washed with PBS and fluorescence was recorded.

### 2.7. Cell exposure for RNA sequencing experiment

HepG2 cells were allowed to grow for 24 h, at a density of 10^6^ cells/ml, in a 25 cm^2^ cell culture flask. Then, the medium was replaced with the fresh cell culture medium containing ZnO NPs and SWCNT. Cells with only a fresh cell culture medium were used as a control sample. Low cytotoxic concentrations were chosen (ZnO NPs: 2 µg/ml, SWCNT: 10 µg/ml) for further transcriptomic analysis at which more than 80% viability was achieved (according to cell viability assay).

### 2.8. RNA purification and library construction: RNA isolation and sequencing

Total RNA was extracted from nanoparticles treated and untreated (control) HepG2 cells using the RNeasy mini kit (Qiagen GMBH, Hilden, Germany) following the manufacturer’s recommendations. The integrity of total RNA was assessed using the Agilent 2100 bioanalyser (Agilent Technologies, Santa Clara, CA, USA) (Figure S1, S2 and S3). The NEBNext® Ultra™ Directional RNA Library Prep Kit (New England Biolabs, MA, USA) was used to prepare the RNA-seq library of an individual sample. The average RIN was >8.0 (Figure S4), and the 1 μg of pooled total RNA was taken for mRNA isolation, fragmentation and priming. Fragmented and primed mRNA was further subjected to the first-strand synthesis in Actinomycin D (Gibco, Life Technologies, CA, USA), followed by second-strand synthesis. The double-stranded cDNA was purified using HighPrep magnetic beads (Magbio Genomics Inc, USA). Purified cDNA was end-repaired, adenylated and ligated to Illumina multiplex barcode adapters. Adapter ligated cDNA was purified using HighPrep beads and was subjected to 12 cycles of Indexing-PCR (37 °C for 15 mins followed by denaturation at 98°C for 30 sec, cycling (98°C for 10sec, 65°C for 75sec) and 65°C for 5mins) to enrich the adapter-ligated fragments. The final PCR product (sequencing library) was purified with HighPrep beads, followed by a library-quality control check. The Illumina-compatible sequencing library was quantified by Qubit fluorometer (Thermo Fisher Scientific, MA, USA), and its fragment size distribution was analysed on Agilent 2200 Tapestation. Samples were then sequenced using the Illumina Genome Analyzer (HiSeq 2500v4 High Output).

### 2.9. Data analysis

FastQC (version 0.11.2) was used for quality control to ensure that the quality value was above Q30. The Human (*Homo sapiens*) genome FASTA file (ftp://ftp.ensembl.org/pub/release-100/fasta/homo_sapiens/dna/) and gene annotation GTF file (Human (GRCh38.p13) assembly; ftp://ftp.ensembl.org/pub/release-100/gtf/homo_sapiens/) were obtained from Ensembl. Although RNA-seq is a popular research tool, there is no gold standard for analysing the data. Therefore, we chose up-to-date open-source tools for mapping, retrieving read counts, and differential analysis among the available tools. We used HISAT2 (version 2.0.3-beta) (Kim et al., 2015) to generate indexes and map reads to the human genome. For assembly, we chose SAMtools (version 1.2) and the “union” mode of HTSeq (version 0.6.1) (Anders et al., 2015), as the gene-level read counts could provide more flexibility in the differential expression analysis. Both HISAT2 and HTSeq analyses were conducted in the Linux operating system (version 2.6.32). Differential expression and statistical analysis were performed using DESeq2 (release 3.3) in R (version 3.2.4). DESeq2 (Love et al., 2014) was chosen as a popular parametric tool that provides a descriptive and continually updated user manual (https://bioconductor.org/packages/release/bioc/vignettes/DESeq2/inst/doc/DESeq2.html).

DESeq2 internally corrects library size, so it is essential to offer un-normalised raw read counts as input. We used variance stabilising transformation to account for differences in sequencing depth. P-values were adjusted for multiple testing using the Benjamini-Hochberg procedure (Benjamini and Hocheberg, 1995). A false discovery rate adjusted p-value (i.e., q-value) < 0.05 was set to select DE genes.

### 2.10. Bioinformatics analysis

The list of differential expression of genes was utilised to observe transcriptional snapshots through bioinformatics analysis. First, the GO and pathway were initially identified through the DAVID analyses using the right-tailed Fisher Exact test and displayed z-scores. Then, the overrepresentation test was performed in PANTHER version 11.1 (http://www.pantherdb.org/, released on October 24, 2016) using the binomial test and Bonferroni correction for multiple testing.

In PANTHER, only pathways and GO terms with fold enrichment > 0.2 and p-value > 0.05 were listed. The pathway enrichment analysis of up and downregulated genes was performed using the R package, cluster profile package (Yu et al., 2012) with a p-value cutoff of ≥ 0.05. All the graphical analyses were performed in the R environment using respective tools. Histograms, density scatter plots, Box plots, Volcano plots and Principal Component Analysis (PCA) were generated using the ggPlot2 package. Finally, all the identified genes were mapped with Uniprot human chromosomal proteome information and parsed to create files appropriately formatted for input to Circos (Krzywinski et al., 2009).

### 2.11. Statistical analysis

The data were analysed using GraphPad Prism 5.01 (GraphPad Software, San Diego, California, USA, https://www.graphpad.com/). The significance of the difference between mean values was calculated by one-way analysis of variance (ANOVA) with Tukey’s posthoc test. The data were represented as mean ± SD, and the value of p < 0.05 was considered statistically significant.

## 3. Results and Discussion

### 3.1. Dose determination and cytotoxicity assessment for nanoparticles treatment

The cytotoxicity assays were performed on HepG2 cells to select the suitable dose and acute exposure for transcriptomics analysis. The aim is to find the non-cytotoxic dose that would better mimic the real-life exposure scenario and predict actual nano-bio interactions. Most of the studies previously performed considered a high dose, acute exposure of NPs considering various animal and cellular systems. These extreme exposures overwhelm the cellular system, consequently leads to necrosis and ultimately death. The cell viability assays (Figure 1) were performed in HepG2 cells exposed to ZnO NPs and SWCNT in a broad concentration (0, 1, 2, 5, 10, 20, 50 µg/ml) range and time (3, 6, 12, 24 h).

**Figure 1:**
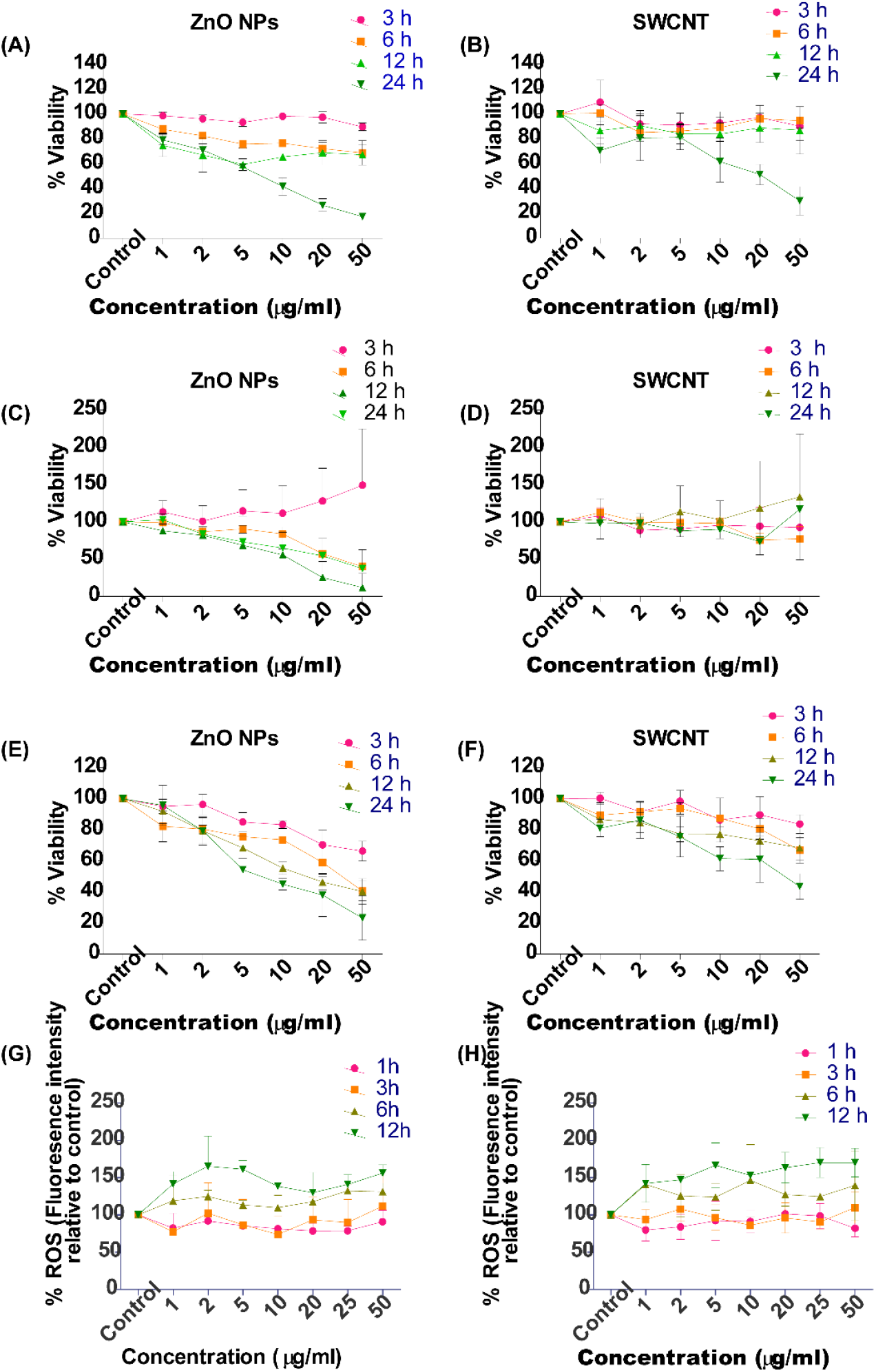
The cell viability and oxidative stress analysis in HepG2 cells exposed to ZnO NPs and SWCNT. (A) MTT, (B) Trypan blue assay and (C) DCFH-DA assay and (D) Nitic oxide assay was performed to assess the cellular viability after exposure of ZnO NPs and SWCNT for 3, 6, 12 and 24 h in a concentration (1, 2, 5, 10, 20, 50 µg/ml) dependent manner. All the assay values were normalised with non-treated control, the viability of which is represented as 100%. Results are shown as mean values ± SD obtained from three independent experiments (n=3). Statistical analysis was performed using two-way ANOVA followed by Bonferroni post-test.

In the MTT assay, significant effects were observed in ZnO NPs (Figure 1A) even at low concentrations than SWCNT (Figure 1B). No significant impact was seen on the cellular viability at 3 h of exposure. However, at 6 h incubation ZnO NPs was able to compromise the cell viability significantly (p<0.05; ∼ 75% cell viability) at a concentration of 5 µg/ml while the same dose of SWCNT was non-toxic. At a dose of 10 µg/ml, significant (p<0.001) diminution of the cell viability of about 70% at 6 h, 60% at 12 h and <50% at 24 h was observed for ZnO NPs.

On the other hand, SWCNT is not causing harm to the HepG2 cells at the same dose even at the exposure of 12 h. The highest ZnO NPs dose, 50 µg/ml, had the prominent (p<0.001) effect with almost 50% diminished viability at 12 h, and 80% loss at 24 h was found, whereas SWCNT at the same dose was observed to be toxic at 24 h only. This observation indicates that ZnO NPs is responsible for causing extensive damage to cells and is more harmful than SWCNT. Therefore, we did not pursue after 50 µg/ml dose. In contrast to MTT, the results of NRU assay (Figure 1) showed significantly (p<0.01) compromised lysosomal integrity at a lower concentration of ZnO NPs (Figure 1C) (2 µg/ml) following exposure to 6 h, but cells maintained the viability by more than 80% as compared to control. With the increasing concentration of ZnO NPs, the viability of the HepG2 cells decreased at 6 h of incubation, whereas the SWCNT (Figure 1D) was not affecting the viability at any of the concentrations. At 12 h of exposure, the viability was significantly (p<0.05) reduced to 40% at 10 µg/ml. More subtle effects were seen at increased doses, i.e. 20 and 50 µg/ml, with significant (p<0.0001) loss of viability to almost 80%.

The assay results of MTT and NRU are quite contradictory because both rely on different mechanisms, therefore, showing other cytotoxicity measurements, majorly in SWCNT. One of the primary reasons could be the potential interaction of the NPs with the assay components that can confound the results obtained (Ong et al., 2014, Breznan et al., 2015). To overcome the issue, we also performed a trypan blue assay (Figure 1E and F) based on the direct dye exclusion from live cells and could eliminate the interference (Hoskins et al., 2012). However, the interaction of NPs with the cells was so strong that even with subsequent washes, they remain adhered with the cells during early incubation (Tsai et al., 2011), but as the cells remain in the presence of NPs, they lose their adherence properties and detach from the cell culture plates specifically during 24 h. This behaviour of cells was observed in the case of ZnO NPs more abruptly than SWCNT, as described previously that carbon nanotubes can act as a scaffold (Rafeeqi and Kaul, 2010). Trypan blue showed concentration and time-dependent manner reduction in cell viability most prevalent at high dose (50 µg/ml) of ZnO NPs (Figure 1E) when exposed for 6 h but as incubation time increases the significant cellular viability was compromised at 5 µg/ml of dose. In the case of SWCNT (Figure 1F), the amount at which cell viability is almost 70% compromised was found to be 10 µg/ml after incubation of 24 h.

We investigated the intracellular production of reactive oxygen species (ROS) as an NPs mechanism to induce cell death. The oxidative stress levels were minimal at the lower incubation exposure period, i.e. 1 h and 3 h, but a slight increase was observed at 6 h at higher doses of >20 µg/ml. In contrast, the exposure of NPs for 12 h exerted the effect at the lower dose of 2 µg/ml in the case of ZnO NPs (Figure 1G). The SWCNT increases ROS level at 6 h of exposure, even at the lowest dose (1 µg/ml) (Figure 1H). Considering the results, 10 μg/ml concentration of SWCNT was chosen as a non-cytotoxic dose showing no significant cell death but indicating some response against the external exposure. Thus, it corresponds to a sub-chronic rather acute toxic response.

### 3.2. Overview of Transcriptional landscape of HepG2 cells responded to ZnO NPs and SWCNT

To elucidate the underlying molecular response of the biological system at a lower dose and short term exposure of NPs, we performed a genome-wide relative transcriptomic investigation in HepG2 cells. The present NGS based comprehensive RNA sequencing study generated about ∼200 million reads at Illumina HiSeq 2500 platform (Figure 2A).

**Figure 2:**
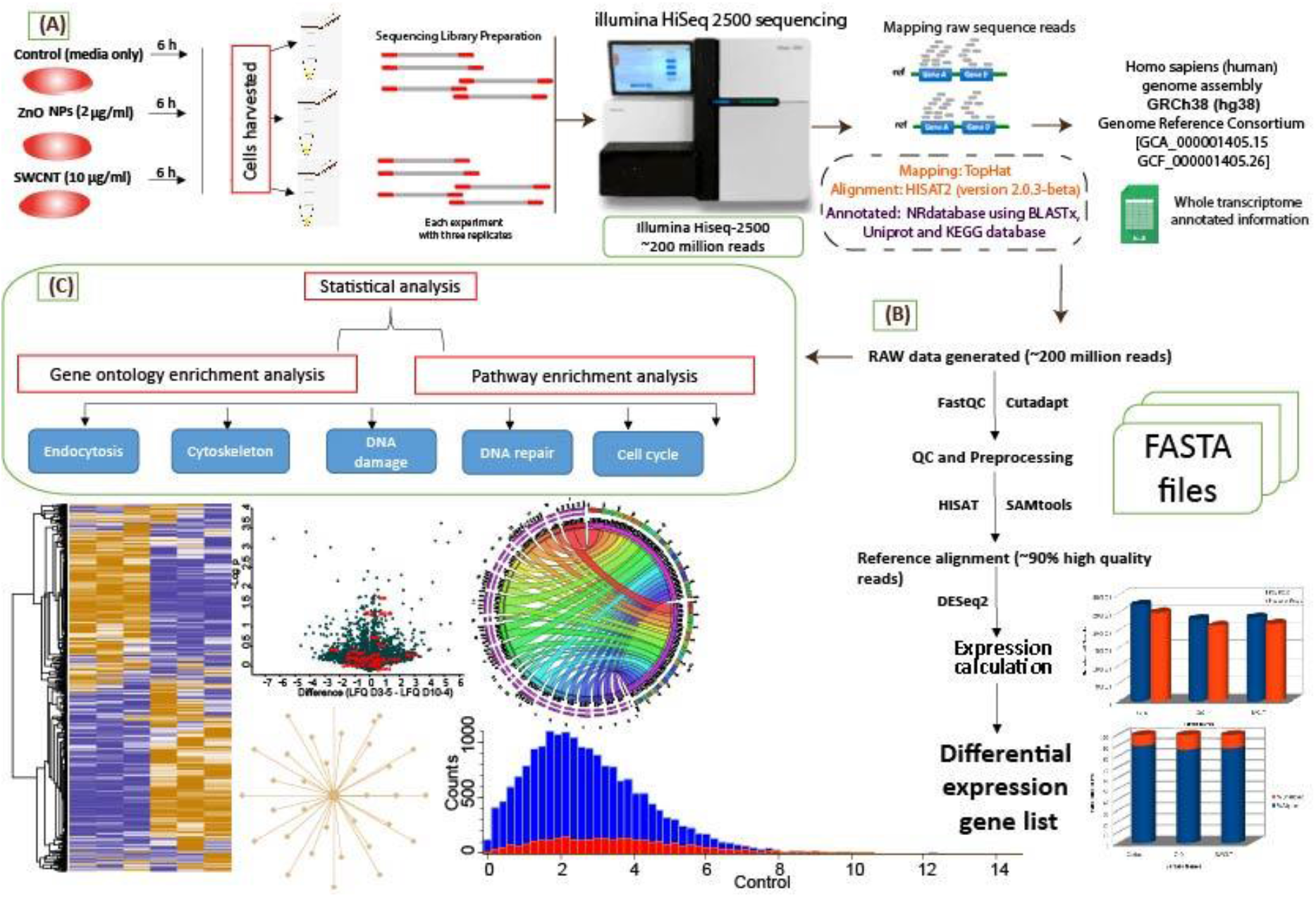
Schematic presentation of complete work flow for the analysis of nanoparticle exposure to HepG2 cell line.

The ample number of reads showed the robustness of the experiment and provided a strong background to identify trustable results. The sequenced basic information was quality checked using FastQC and pre-processing by Cutadapt, which includes removing adapter sequences and low-quality bases (<q30). The high-quality data were then aligned to the reference genome of *Homo sapiens* GRCh38 and classified into aligned and unaligned reads using fast and sensitive spliced alignment programme HISAT. Approximately 90% of sequence reads were aligned to GRCh38genome (Figure S5) signifies superiority of reads and excellent mapping efficiency. After the normalisation, a total of 150 million high-quality adapter free reads was obtained that were utilised for further analysis (Figure 2B). In this way, the 50 million reads per sample were retained for downstream analysis. Precisely, the obtained million reads are 54.323 for control, 46.249 for ZnO2 and 47.337 in the case of SWCNT10. The transcript abundance was estimated in the form of FPKM values for measuring gene/transcript expression. The comparisons were made between the control, and NPs treated samples which revealed differential expression (DE) of 11,027, expressed genes; 5,474 upregulated (p ≤0.05; 277), 5,553 downregulated (p ≤0.05; 215), genes in ZnO NPs and 8,099; 4,287 upregulated (p ≤0.05; 331), 3,812 downregulated (p ≤0.05; 146) genes in SWCNT. The transcripts with log2 expression value ≥1.5 and log2 ≤0.66 were considered as upregulated and downregulated, respectively.

Furthermore, the data was scrutinised, and we considered ≥ 3 folds up and downregulated genes (Table S1) for the further analysis process. This strategy was adopted to provide more stringent results and essential information about highly regulated genes. As we have considered acute exposure conditions of NPs, we hypothesised that the sudden or rapid expression of genes during the early exposure would provide the actual snapshot of NPs effect on the cellular system. With the fold change cut off, a total of 1762 upregulated and ZnO2 NPs deregulated 1163 downregulated genes and for the treatment of SWCNT10, 1195 upregulated, and 1124 downregulated genes were found (Table S1).

The total differential expressed transcriptome classification revealed a high number of contributing mRNAs from chromosomes 2, 17 and 19. In contrast, the minor transcripts were expressed by 13, 18, 21, and Y chromosomes (Figure 3A). The normalised data represent the cumulative equal abundance (Figure 3B) and detailed the normally distributed data (Figure 3C). The coefficient of correlation for FPKM values is more than 0.98 (Figure 3D and E).

**Figure 3:**
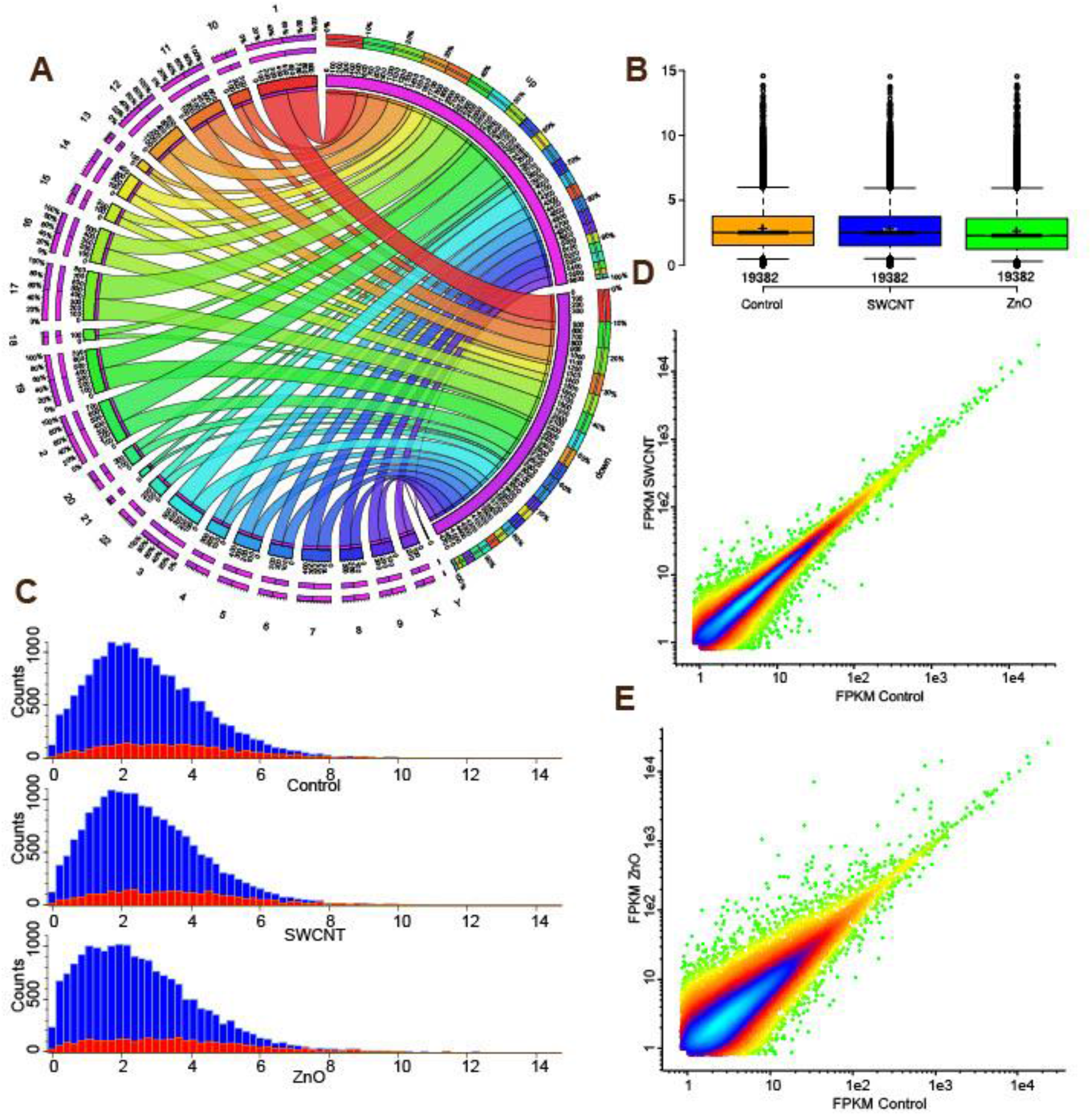

The quality check showed the best segregation of the nanoparticles treated transcriptome (Figure 4A). We highlighted the specific cell cycle-associated and DNA damage proteins in the rank plot (Figure 4B and C).

**Figure 4:**
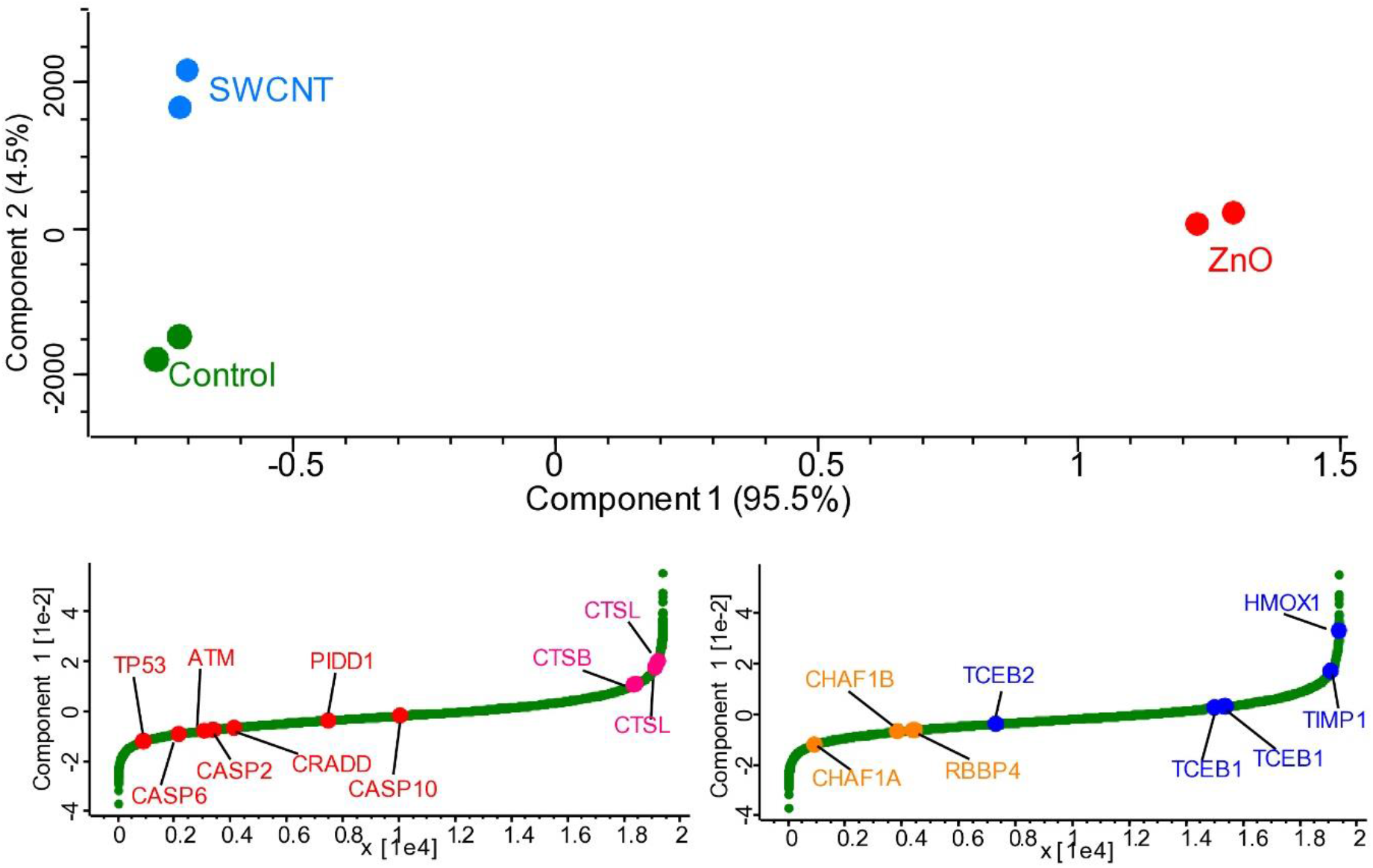
Principle Component analysis. Complete PCA analysis and rank plot of differentially abundant transcripts.

### 3.3. Ontological classification of genes altered from ZnO NPs and SWCNT treatment

Using several ontological databases, differentially expressed genes for both NPs were distributed among various gene ontology (GO) categories (Figure 2C) such as biological process (BP), cellular component (CC), molecular function (MF), and pathways to answer the critical biological questions of what, when, and why these genes are expressed in response to NPs. We selected these ontological terms based on their higher significance of individual p≤0.05 with multiple hypothesis 2 post-test corrected p values (p≤0.05) Bonferroni and (p≤0.05) Benjamini corrected followed with the FDR less than 0.05. In this way, the deciphered key functions better interrogate the interaction of NPs with HepG2 liver cells. Even after filtering to multiple tight parameters, high degree of consistency between the ontologies were found related to endocytosis (GO:0006897; ZnO2 p-value 1.55E-05, SWCNT10 p-value 1.11E-04), ubiquitination (GO:0042787; ZnO2 p-value 5.49E-04, SWCNT10 p-value 0.027198), cell-cell adhesion (GO:0098609; ZnO2 p-value 8.89E-09, SWCNT10 p-value 0.011429), cellular response to DNA damage stimulus (GO:0006974; ZnO2 p-value, 1.83E-04, SWCNT10 p-value 0.014698137), DNA repair (GO:0006281; ZnO2 p-value 8.81E-04, SWCNT10 p-value 21.33E-06) and cell migration (GO:0016477; ZnO2 p-value 0.06196377, SWCNT10 p-value 0.018241486) were significantly over enriched for both the NPs groups.

Furthermore, we also identified a considerable number of genes scattered in different locations in a cell also obtained through cellular component category genes involved in the cytoplasm (GO:0005737; ZnO2 p-value 4.17E-22, SWCNT10 p-value 1.43E-18), nucleus (GO:005634; ZnO2 p-value 2.93E-14 SWCNT10 p-value 1.08E-12), nucleoplasm (GO:0005654; ZnO2 p-value 1.09E-14, SWCNT10 p-value 2.86E-14), cell-cell adherens junction (GO:0005913; ZnO2 p-value 1.18E-09, SWCNT10 p-value 0.002373005), focal adhesion (GO:0005925; ZnO2 p-value 3.12E-06, SWCNT10 p-value 6.50E-04), and nuclear matrix (GO:0016363; ZnO2 p-value 0.001027326, SWCNT10 p-value 9.64E-04) were most significantly counted. In terms of molecular function, the most significant terms were related to protein binding (GO:0005515), zinc ion binding (GO:0008270), ATP binding (GO:0005524) (Table S2). Based on the categorisation of the GO, we distinguish six reflex processes of HepG2 cells, including DNA damage, DNA repair, cell cycle, cytoskeleton, transcriptional changes, ubiquitin-mediated proteasomal degradation of proteins. The elaborated list (with FPKM values and Fold change) of common and unique genes for ZnO NPs and SWCNT, considered for the subsequent preparation of the Venn diagram, is provided in the supplementary TableS4. We have observed similar and unique processes against the NPs insult, but we are interested in looking closely at the processes and their associated genes that the two different types of NPs have commonly perturbed.

### 3.4. The interplay of NPs internalisation and subsequent changes in the cytoskeletal structure of the cells

The process of endocytosis considers the central gateway through the cell. Various materials can also exploit the process of endocytosis to gain entry inside the cells (Oh and Park et al., 2014). However, the NPs must be internalised by the cells to carry out their functions or, in other words, to regulate cellular processes. Therefore, unravelling the role of the internalisation mechanism opens up new avenues in the design and development of novel nanocarrier systems and their potential effects. In this frame, our RNA sequencing data suggested a significant enrichment of endocytosis (GO:0006897) (Table S3) process for the log_2_ fold change >3 deregulated genes in case of both the ZnO2 (p-value 1.55E-05) NPs and SWCNT10 (p-value 1.11E-04). Strikingly, 25% of the common upregulated and 21% downregulated transcripts were observed by the Venn comparisons for the process of endocytosis (Figure 5A), giving clues of the potential intracellular trafficking network of NPs.

**Figure 5:**
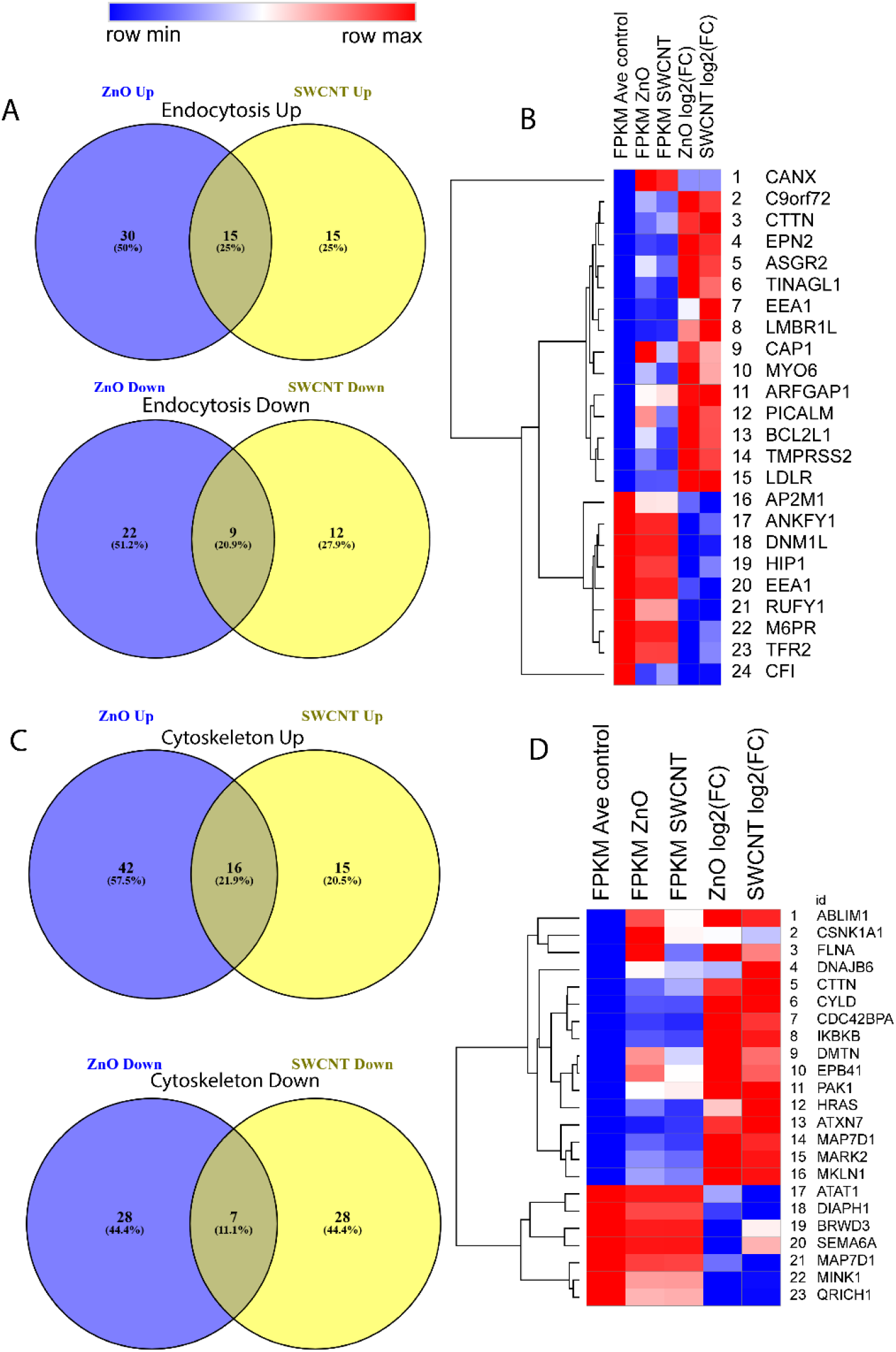
Summary of ZnO NPs and SWCNT perturbed gene detected by RNA-sequencing, distributed for Endocytosis and Cell cytoskeleton related biological process. Venn diagram of differentially expressed genes representing overlapped and NPs specific upregulated and downregulated genes with FC ≥3 in the biological process categories (A) Endocytosis and (C) Cell cytoskeleton. (B, D) Showing heatmap of the genes detected in the overlap of the Venn diagrams changed by both the NPs. Columns are ordered by FPKM values of control, ZnO NPs, SWCNT and calculated values of log2 fold change. Blue to red denotes low to high expression as shown on the scale. The common upregulated genes representative of the endocytosis process includes *CANX, CTTN, EPN2, LMBR1L, CAP1, MYO6, ARFGAP1, PICALM, BCL2L1, TMPRSS2* and *LDLR*.

Notably, subsequent heatmap analysis (Figure 5B) showed a similar level of this transcript abundance based on their FPKM values regulated in response to ZnO2 NPs and SWCNT10. Before the NPs reach the exterior membranes, they must interact with the microenvironment around the target cells. The dynamic physiochemical interactions play a significant role at the nano-bio interface that governs the actual effect of NPs on a cell (Nel et al., 2009). When the NPs first encounter the extracellular components, they contact various biomolecules such as proteins (or biomolecules secreted by cells in the medium) and form a protein corona that is majorly responsible for changing the physicochemical composition of NPs. These newly acquired properties have a profound effect on the cell-NPs interactions (Francia et al., 2019) wherein, they can act as a ligand to be engaged with cell surface receptors to enhance the uptake of NPs (Lara et al., 2017). This could be explained, at least partly, by the high upregulation of the LDLR gene (Low-density lipoprotein receptor) (ZnO2 FC: 12.46, SWCNT10 FC: 12.49), giving us a clue about recognition of protein corona by cell-surface LDL receptor. In another way, it can be correlated to the secretion of Apolipoprotein by HepG2 cells and its interaction with NPs as one of the constituents of protein corona and further interaction with the LDL receptor mediating the receptor-mediated endocytosis of NPs.

Cellular internalisation can initiate various processes, including the formation of early endosomes and sort of the outside material, just as in the case of ZnO2 NPs and SWCNT10 suggested by the significantly enriched CC GO term early endosome (GO:0005769: ZnO2 p-value 3.57E-04, SWCNT10 p-value 0.002291876) (Table S3). As previously defined, much cytoskeletal protein machinery plays a significant role during the process of endocytosis. Most importantly, actin polymerisation provides a sufficient force for membrane invagination and resulting in vesicle budding. It is anticipated that dynamic changes in the actin structures are a crucial step in the process of clathrin-mediated endocytosis that ultimately provides enough force for membrane bending (Skruzny et al., 2012). It is also explained by significant GO enrichment of actin cytoskeleton reorganisation (GO:0031532) in our transcriptomics data with subsequent cellular component category including actin cytoskeleton (GO:0015629: ZnO2 p-value 0.001249031, SWCNT10 p-value 0.020259711) and focal adhesion (GO:0005925: ZnO2 p-value 3.12E-06, SWCNT10 p-value 6.50E-04) for both the NPs (Table S3). The upregulation of CTTN, cortactin protein (ZnO2 FC: 6.5, SWCNT10 FC: 7.2) (Figure 5C and D), responsible for vesicle scission from the plasma membrane (Cao et al., 2003).

Consistent with these observations, we can say that NPs internalisation is correlated with the altered cytoskeletal structure and cells ability to sense the changing biophysical properties of extracellular matrix (ECM). On the other hand, altering ECM due to NPs internalisation alters other critical cellular functions such as proliferation, migration, cell membrane structure, and organisation (Panzetta et al., 2107). This is evidenced in our transcriptomics dataset where the major BP category comprised significantly cell-cell adhesion (GO:0098609: ZnO2 p-value 8.89E-09, SWCNT10 p-value 0.011428656) and cell migration (GO:0016477: ZnO2 p-value 0.06196377, SWCNT10 0.018241486).

It was also mentioned that NPs could gain access to the cellular microenvironment through non-specific interactions (Behzadi et al., 2017). Cells from mechanically active environments can couple mixed signals from forces applied through β-integrins to upregulate the production of cytoprotective cytoskeletal proteins, typified by filamin A (FLNA) (ZnO2 FC: 8.2, SWCNT10 FC: 6.2) (Figure 5D) (D’Addario et al., 2001).

### 3.5. Impact of Nanoparticles on the DNA Damage and Repair Related Genes

Cells maintain highly complex signalling processes and are programmed to sense the damage. Thus, it regulates multifaceted responses to maintain the cellular physiology against external stimuli. Previously many studies have identified the DNA damaging potential of NPs at higher concentrations. For instance, Sharma et al observed that the ZnO NPs could cause oxidative stress-mediated DNA damage in HepG2 cells (Sharma et al., 2012). Similarly, Chk1-dependent G2/M, DNA damage checkpoint signalling pathway, activated in response to silica NPs at 100 µg/ml, resulted in cell cycle arrest due to oxidative DNA damage (Duan et al., 2013). One of the studies conducted in our lab observed potential DNA damage in buffalo sperms when exposed to lower concentrations of TiO_2_ NPs (Pawar and Kaul, 2014). In the present study, we observed DNA damage even at the low, non-cytotoxic doses of NPs (as described by cytotoxicity analysis, Figure 1A-H) and interestingly found that both NPs exert similar responses at the transcriptomic level (Table S3). The comparison can be easily made in Venn’s diagram analysis prepared by genes of enriched DNA damage category that spans 21% upregulated genes, and 7% downregulated genes (Figure 6A) altered by both the NPs.

**Figure 6:**
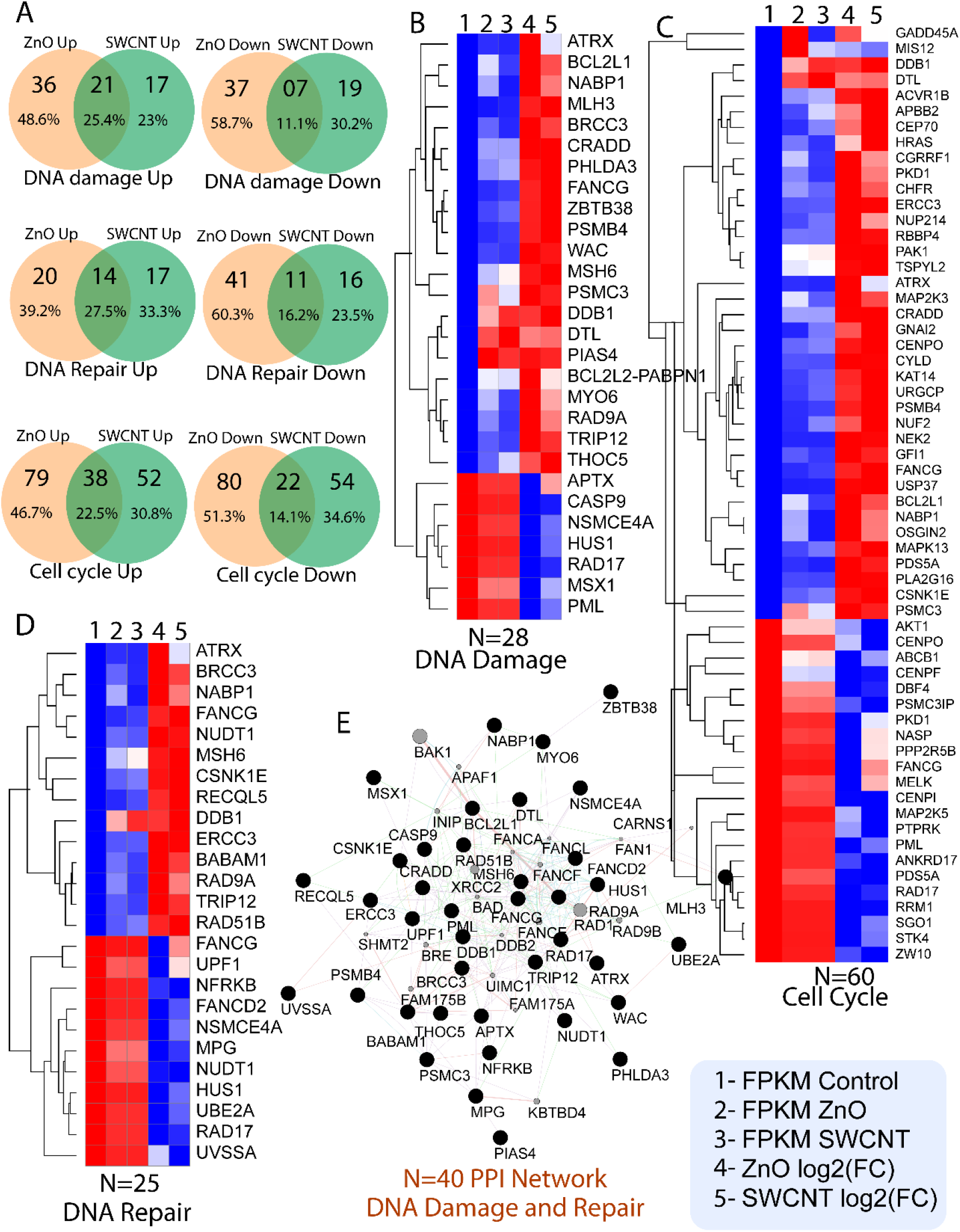
Summary of ZnO NPs and SWCNT perturbed gene detected by RNA-sequencing, distributed for DNA damage, repair and cell cycle-related biological process. (A) Venn diagram of differentially expressed genes representing overlapped and NPs specific upregulated and downregulated genes with FC ≥3 in the biological process categories DNA damage and DNA repair, and Cell cycle. (B, C, D) Showing heatmap of the genes detected in the overlap of the Venn diagrams changed by both the NPs. Columns are ordered by FPKM values of control, ZnO NPs, SWCNT and calculated values of log2 fold change. Blue to red denotes low to high expression as shown on the scale. (E) The PPI network of DNA damage and DNA repair.

A similar observation was captured previously, where similar expression patterns for different types of NPs (TiO2 NPs and MWCNT) during 4 h of incubation were induced in the cell as a general mechanism for toxicity (Tilton et al., 2014).

Venn diagram also represents 48.6% genes for ZnO2 NPs, 23% genes for SWCNT10 to be upregulated, and 58.7% for ZnO2 NPs and 30.2% downregulated genes SWCNT10 in the DNA damage process (Figure 6A). The presence of multiple GO terms including DNA repair (GO:0006281; ZnO2 p-value 8.81E-04, SWCNT10 p-value 1.33E-06), cellular response to DNA damage stimulus (GO:0006974; ZnO2 p-value 1.83E-04, SWCNT10 p-value 0.014698137) and mitotic cell cycle checkpoint (GO:0007093; ZnO2 p-value 0.016635545, SWCNT10 p-value 0.001777846) were shown to be significantly altered (Table S3) and proves that NPs treatment resulted in the significant disturbance in these processes (Figure 6B-E). Hierarchical clustering of the genes enriched for DNA damage and repair (Figure 6B and D) reveals potential deregulated transcripts in response to NPs exposure. These enriched pathways propose that DNA damage be persuaded in HepG2 after exposure to ZnO2 NPs and SWCNT10.

Generally, the DNA repair system gets activated once the cells experience damage induced by NPs to minimise the negative impact of the injury on cell proliferation (Carriere et al., 2016). The checkpoint network must sense the damage and promptly spread such signals to reach the downstream cellular effector proteins. The perfect correlation among those genes was observed in the similarity analysis, where malfunction of Rad17 (ZnO2 FC: -11.3, SWCNT10 FC: -10.3), a critical signalling node of DNA damage signalling, has been linked to various cancer statuses. The upregulation of RAD related proteins provides a high-fidelity tolerance mechanism against DNA damage stress. The upregulated gene RAD51B (ZnO2 FC: 5.6, SWCNT10 FC: 6.11) takes part in homologous recombination repair and influence the cell cycle progression against DNA damage (Lee et al., 2014) (Figure 6E). We also observed that upon stimulation to NPs, specific categories of genes were upregulated that actively takes part in DNA repair pathways such as DDB1 (ZnO2 FC: 3.4, SWCNT FC: 3.8). DDB1 forms a heterodimer with DDB2, subsequently with CUL4A E3 ligase that actively mediates the ubiquitination of target proteins such as histones and ensures efficient activation of nucleotide excision repair at the DNA lesion site (Li et al., 2006). This CUL4A E3 ligase complex interacts with another upregulated gene DTL (ZnO2 FC: 3.2, SWCNT10 FC: 3.4) is necessary for early G2/M checkpoint (Sansam et al., 2006) and protecting HepG2 cells.

A regulatory kinase gene CSNK1E significantly overexpressed (ZnO2 FC: 14, SWCNT10 FC: 14.4) responsible for phosphorylating p53 under stress conditions, consequently weakening its interaction with its cellular counterpart MDM2 and activates it (p53 phosphorylation, phosphorylation p53 by CK1, CK1 p53 threonine kinase). But TP53BP1 is downregulated, and MDM2 is upregulated in our case. This is explained as the levels of casein kinase is very high (14 FC), and its starts are phosphorylating p53 to activate. It also regulates the beta-catenine dependent WNT pathway positively by phosphorylating and stimulating the activity of the dishevelled protein (Klimowski et al., 2006). Another gene from serine/threonine kinase family PLK3 is involved in the cell cycle progression, apoptosis (Wang et al., 2002), and DNA damage was downregulated (Xie et al., 2001). In addition, it phosphorylates and regulates the number of other effector molecules such as Bcl-xL (Wang et al., 2011), Chk2 (Bahassi et al., 2006), p53 (Bahassi et al., 2002) and ATF2 (Wang et al., 2011). In particular, some of the phosphorylation targets of PLK3, including Bcl-xL and ATF2, are upregulated by NPs, but others (Chk2) are downregulating. The gene PLK3 also cause degradation of wee1 kinase (downregulated in the cell cycle), a negative regulator of the cell cycle involved in re-entry into cell cycle progression (Figure 6C).

The NPs treated cells show the downregulation of some cell cycle checkpoint protein such as RAD17 (rad17 cell cycle checkpoint). It may be due to proteolysis by cdh1/APC, resulting in the termination of checkpoint signalling and recovery from genotoxic stress. In summary, our low dose transcriptomics analysis proves that ZnO2 NPs and SWCNT10 could induce DNA damage, oxidative stress and affect the HepG2 cells. Our study provides a comprehensive analysis of how exposure to ZnO2 NPs and SWCNT10 on HepG2 cells could affect its molecular landscape and whether these changes are beneficial or detrimental for the HepG2 cells (Figure 6E).

### 3.6. Transcriptional changes in response to NPs

It is crucial to know how the NPs regulate various genes’ spatial and temporal expression by altering the transcriptional machinery for mRNA synthesis (Figure 7A).

**Figure 7:**
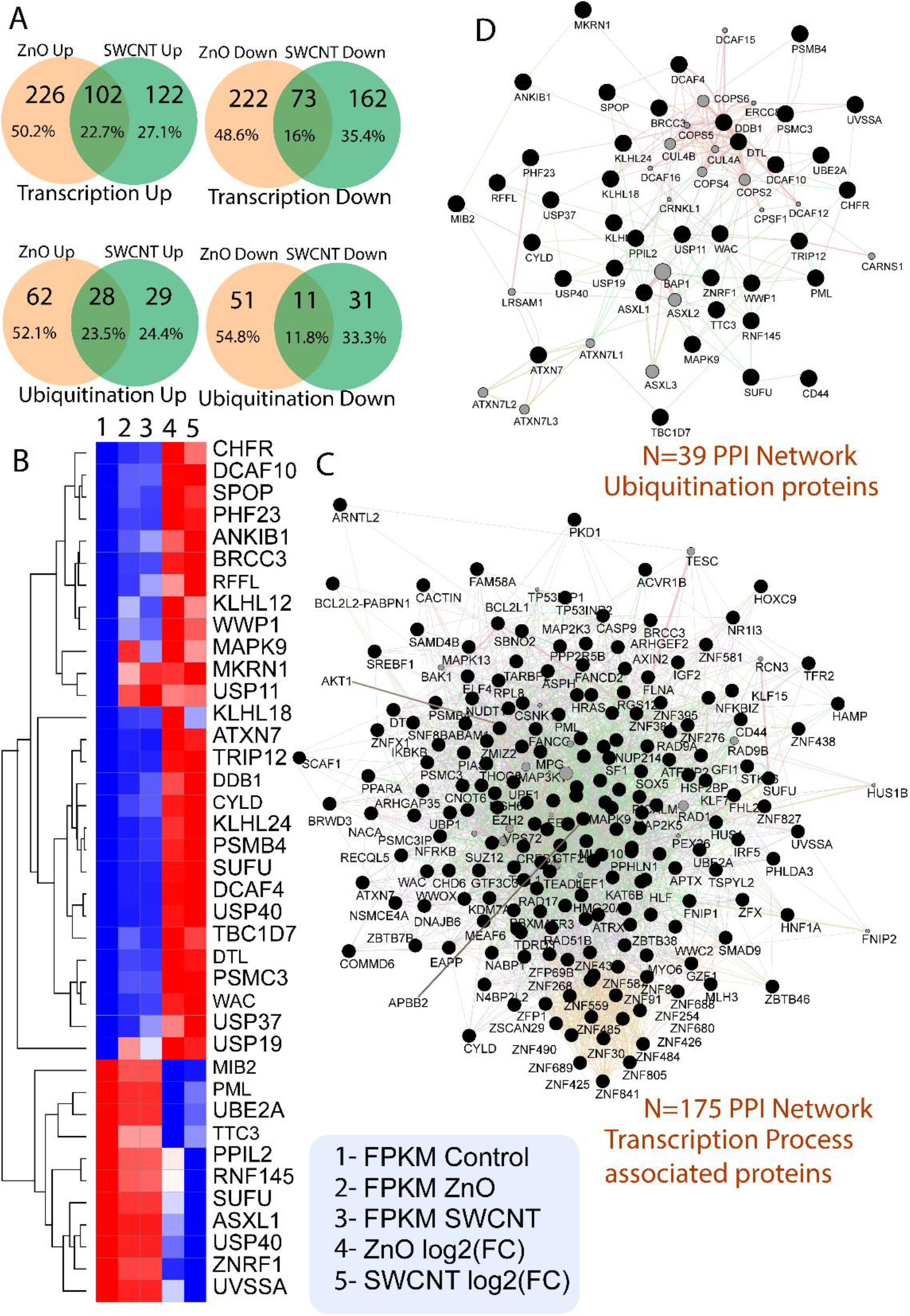
(A) Venn diagram of differentially expressed genes representing overlapped and NPs specific upregulated and downregulated genes with FC ≥3 in the transcription and ubiquitination related biological process categories. (B) Showing heatmap of the ubiquitination genes detected in the overlap of the Venn diagrams changed by both the NPs. Columns are ordered by FPKM values of control, ZnO NPs, SWCNT and calculated values of log2 fold change. Blue to red denotes low to high expression as shown on the scale. (C) The PPI network of transcription related biological process. (D) The PPI network of ubiquitination related transcripts.

In this context, the poly (A) tail is the major determining factor for the turnover rates of mRNAs under different cellular conditions. We found that the gene expression changes resulted from NPs treatment in mRNA turnover through major identified GO terms (Table S2). This includes nuclear-transcribed mRNA poly (A) tail shortening (GO:0000289; ZnO2 p-value 1.41E-05), regulation of mRNA stability (GO:0043488, ZnO2 p-value 0.0010776), regulation of transcription DNA templated (GO:0006355, ZnO2 p-value 0.004395587), regulation of DNA-templated transcription in response to stress (GO:0043620; ZnO2 p-value 0.010266263), positive regulation of cytoplasmic mRNA processing body assembly (GO:0010606; ZnO2 p-value 0.010266263), negative regulation of transcription from RNA polymerase II promoter (GO:0000122; ZnO2 p-value 0.024209881), regulation of translational initiation (GO:0006446; ZnO2 p-value 0.030648445), and histone deacetylation (GO:0016575; ZnO2 p-value 0.039595812) specifically observed to be upregulated in ZnO2 NPs. Similarly in case of SWCNT the enriched terms related to transcription includes regulation of sequence-specific DNA binding transcription factor activity (GO:0051090; SWCNT10 p-value 3.59E-04), covalent chromatin modification (GO:0016569; SWCNT10 p-value 0.001780004), transcription, DNA-templated (GO:0006351; SWCNT10 p-value 0.004554925), chromatin remodelling (GO:0006338; SWCNT10 p-value 0.009144073), posttranscriptional regulation of gene expression (GO:0010608; SWCNT10 p-value 0.010799581), regulation of mRNA stability (GO:0043488; SWCNT10 p-value 0.031509087), and histone deacetylation (GO:0016575; SWCNT10 0.044248664).

Ribonuclease specific to Poly (A) tail, PARN (ZnO2 FC: 4.1, SWCNT10 FC: 5.2) was upregulated that is responsible for efficient degradation of mRNA poly (A) tails and found that PARN is also a crucial player, accountable for regulating the dynamics of poly (A) during the DNA damage processes (Cevher et al., 2010). Therefore, our results indicate the silencing or degradation of aberrant mRNAs during the stress condition. In the same context, the upregulation of major subunits of the CCR4-NOT complex (major deadenylase), i.e., CNOT1 (ZnO2 FC: 3.05, SWCNT10 FC: 2.09) and CNOT6 (ZnO2 FC: 7.2, SWCNT10 FC: 9.57) was also observed. CNOT1 maintains cell viability by maintaining the levels of translatable mRNA that, in turn, prevents the overproduction of proteins and ultimately prevents the unfolded protein response (Ito et al., 2011). CNOT6 (ZnO2 FC: 7.2, SWCNT10 FC: 9.57) also contains cell senescence and is vital for efficient cell proliferation (Mittal et al., 2011). Furthermore, we observed that NPs could alter the expression of genes related to chromatin remodelling (GO:0006338) that controls epigenetic mechanisms in the cell (Sierra et al., 2016).

The results of toxicity assays showed that NPs are responsible for DNA damage and proved in transcriptomics results discussed in the previous section. To counteract the defence mechanism, our data show that cells upregulate various chromatin remodellers such as INO80 (ZnO2 FC: 3.4, SWCNT10 FC: 3.5) to modify the nucleosomes in such a way that different repair proteins can gain access to the damaged sites on the DNA (Attikum et al., 2004 6; Morrison et al., 2004). Thus, NPs alter chromatin-based transcriptional machinery to regulate the cellular processes under changing cellular environments. We found AJUBA (ZP 3.6, SM 3.4), which is responsible for HDAC (ZnO2 FC: 2.3 SWCNT10 FC: 1.6) dependent corepressor activity and negative regulator of the Hippo signalling pathway (Ayyanathan et al., 2007; Thakur et al., 2010). AJUBA is also identified to regulate cell-cell adhesion formation and affect cell growth and fate (Marie et al., 2003).

Another interesting finding is the downregulation of the Hepatocyte nuclear factor 1A (HNF1A; ZnO2 FC: -11.8, SWCNT10 FC: -11.5) in both the NPs treatment. It is a major hepatic transcription factor responsible for regulating vital processes, majorly involved in glucose, amino acids, lipid, homeostasis, detoxification, bile acids and cholesterol metabolism. Also, the mutation in HNF1A causes maturity-onset diabetes of the young type 3 (MOYD3), characterised by impaired insulin secretion. The HNF1A knockdown study showed elevated genes related to proliferation and cell cycle control (Bonzo et al., 2012).

In addition, loss of HNF1A activity in liver adenocarcinoma resulted in increased cellular proliferation with an increase in glycolysis, lipogenesis, and other macromolecules that further advantage the growth (Wang et al., 2019). However, in our transcriptomics data, we not only found downregulation of HNF1A (ZnO2 FC: -11.8, SWCNT10 FC: -11.5) but also for other concordant genes, including hepatocyte growth factor activator (HGFAC, ZnO FC: -2.7, SWCNT10 FC: -5) and insulin growth factor 2 (IGF2, ZnO2 FC: -12.9, SWCNT10 FC: -8.7), therefore, resulted in the restricted aberrant cellular proliferation upon NPs treatment. Thus, we evident at the molecular level that a low dose of NPs imposes a negative effect on the HepG2 cells, but the regulatory machinery of cells described above functions in a coordinated manner to protect from the toxic insult and enhance the survival rate.

The highly upregulated gene in the category related to transcription (GO:0006351) is FHL2 (ZnO2 FC: 15.5, SWCNT10 FC: 13.3) that exerts its effect on transcription by showing its inhibitory impact on E4F1, a cell cycle repressor (Paul et al., 2006). The major suppressor KAT6B (ZnO2 FC: 13.5, SWCNT10 FC: 9.2) gene involved in dynamics of histone H3 acetylation is highly expressed in response to NPs that can be a positive regulator of transcription (Champagne et al., 1999). The upregulation of major gene CACTIN (ZnO2 FC: 12.7, SWCNT10 FC: 12.3) suggested TLR mediated NF-kB signalling (Atzie et al., 2010) prevents the inflammatory response during low dose exposure of NPs. The nuclear protein peripherin (PPHL1, ZnO2 FC: 12.28, SWCNT10 FC: 11.78) arrests the cell cycle at S-phase by downregulating the expression of a vital initiation factor of replication cdc7 (Kurita et al., 2004).

Serine/threonine kinases are also found to be activated observed in gene ontology category of molecular function (ZnO2 p-value 0.025, SWCNT10 p-value 1.18E-04; TableS3) that includes MAPK13 (ZnO2 FC: 11.93, SWCNT10 FC: 8.27) phosphorylates variety of proteins and mediates their degradation through proteasomal pathway in stressed conditions (Pani et al., 2008). Some other important kinases in our data includes ACVR1B (ZnO2 FC: 6.59, SWCNT10 FC: 6.9), CDC42BPA (ZnO2 FC: 6.49, SWCNT10 FC: 5.83) CSNK1A1 (ZnO2 FC: 4.62, SWCNT10 FC: 4.25) CSNK1E (ZnO2 FC: 14, SWCNT10 FC: 14.48), LIMK2 (ZnO2 FC: 11.21, SWCNT10 FC: 11.17), MAP2K3 (ZnO2 FC: 8.99, SWCNT10 FC: 7.68), MAPK13 (ZnO2 FC: 11.93, SWCNT10 FC: 12.7), MAPK3 (ZnO2 FC: 6.64, SWCNT10 FC: 7.4), MARK2 (ZnO2 FC: 9.94, SWCNT10 FC: 9.5), MAST2 (ZnO2 FC: 8.75, SWCNT10 FC: 9.3) NEK2 (ZnO2 FC: 7.5, SWCNT10 FC: 7.4), PAK1 (ZnO2 FC: 3.68, SWCNT10 FC: 3.69), PKN1 (ZnO2 FC: 7.9, SWCNT10 FC: 3.2), RIOK3 (ZnO2 FC: 11.10, SWCNT10 FC: 12.86) (Table S1). Pathway analysis also revealed similar pathways induced by both the NPs that included focal adhesion, MAPK signalling and TNF signalling pathway (TableS2). Overall, these finding suggest that ZnO2NPs and SWCNT10 significantly affect the transcriptomic profile of HepG2 cells, leading to measurable alternation in behaviour.

### 3.7. Ubiquitin mediated proteasomal degradation of proteins

The NPs shows the concentration-dependent lethality to the cell; at low concentration, it shows disruption in the cellular machinery by interacting with various biomolecules and proteins inside the cell. The cell always tries to maintain its appropriate environment by targeting misfolded and proteins of DNA repair, apoptosis and many other cellular events. NPs enter inside the cells and interact with various molecules, including proteins, and affect their functioning. Accordingly, the upregulation of different ubiquitin-proteasome genes was observed in response to both the NPs. This activation suggests that NPs are causing harm to the proteins due to which cells come into play to activate its degradative machinery to maintain homeostasis. In other perspectives, this ubiquitination machinery is required for activating and regulating various signalling cascades in a nonproteolytic manner.

The selective proteolysis of truncated proteins is an essential requirement to maintain the cellular homeostasis mainly regulated by coordinating the ubiquitin-proteasome system (UPS). The ubiquitin is the posttranslational modification that particularly targets the modified proteins to proteasomal complex. Our dataset identified ubiquitination as the highly enriched top ten processes in the represented DEGs (Table S2). The transcriptomes’ abundances were elevated greater than log_2_ 3.0 fold observed for 24.4% and 52.1% genes in the transcriptomes in presence with SWCNT10, and ZnO2 NPs, respectively (Figure 7A). The GO functional annotation highlights ubiquitination process for the UPS (GO:0070536) for SWCNT10 (*P*-value 0.038), ZnO2 NPs (*P*-value 0.0024), (Table S2).

Also, the proteasome subunits identified were found elevated (Table S5; Figure 7B and C). The previous report also identified the increased levels potentially related to proteasome of 6 transcripts, *PSMB4, PSMC6, PSMA6, PSMC3, PSMD2, and PSMA7*, elevated by more than 3 log2 fold in response to ZnO2 NPs (Braga et al., 2014). The enrichment of ubiquitination processes describes the previously reported DNA replication, transcription and cellular iron ion homeostasis were significantly downregulated because NPs increase the protein damage and target them for degradation and suggesting that exposure to NPs treatment could lead to DNA damage and cells are trying to cope up with the damage by activating repair mechanism (Figure 7 and Table S2). The stability in the cell was maintained through antagonising ubiquitination and deubiquitinating enzymes (DUBs) while retaining the structure and function of proteins, protein turnover, DNA damage response and several other signalling pathways. (He et al., 2016, Bhattacharyya et al., 2014). We found that the NPs show concentration-dependent lethality to the cell (Figure 1); in the low concentration, it shows disruption in the cellular machinery by interacting with various biomolecules and proteins inside the cell. However, the HepG2 cell maintains its appropriate environment by targeting misfolded as well as proteins of DNA repair (GO:0006281; ZnO2 p-value 8.81E-04, SWCNT10 p-value 1.33E-06), apoptosis (GO:0042981; ZnO2 p-value 0.042, SWCNT10 p-value 0.001) and many other cellular events as identified in the GO enrichment process.

Previously, it reported that entry of the NPs (SWCNT and ZnO) inside the cells initiates multiple pathways through activating and deactivating various proteins and affects their functioning (Yadav et al., 2021, Mukherjee et al., 2018, Wilhelmi et al., 2013, Wang et al., 2020, Ndika et al., 2018). Accordingly, the upregulation of various ubiquitin-proteasome genes was observed in response to both the NPs (Figure 7D). This activation suggests that NPs are causing harm to the proteins due to which cells come into play to activate its degradative machinery to maintain homeostasis. In other perspectives, this ubiquitination machinery is required for activating and regulating various signalling cascades in a nonproteolytic manner. While many transcriptome profiling studies have revealed differentially regulated genes encoding proteolysis-related functions, only minor proportions of these transcripts encoded proteasome subunits. Therefore, this concerted elevation of proteasome subunit transcripts is not a general stress response and may reflect the specific biological effects of these NP types.

Furthermore, our data suggest that NP induce changes in the ubiquitin-proteasome system. We postulated the schematic presentation of the NPs affecting the cellular process (Figure 8).

**Figure 8:**
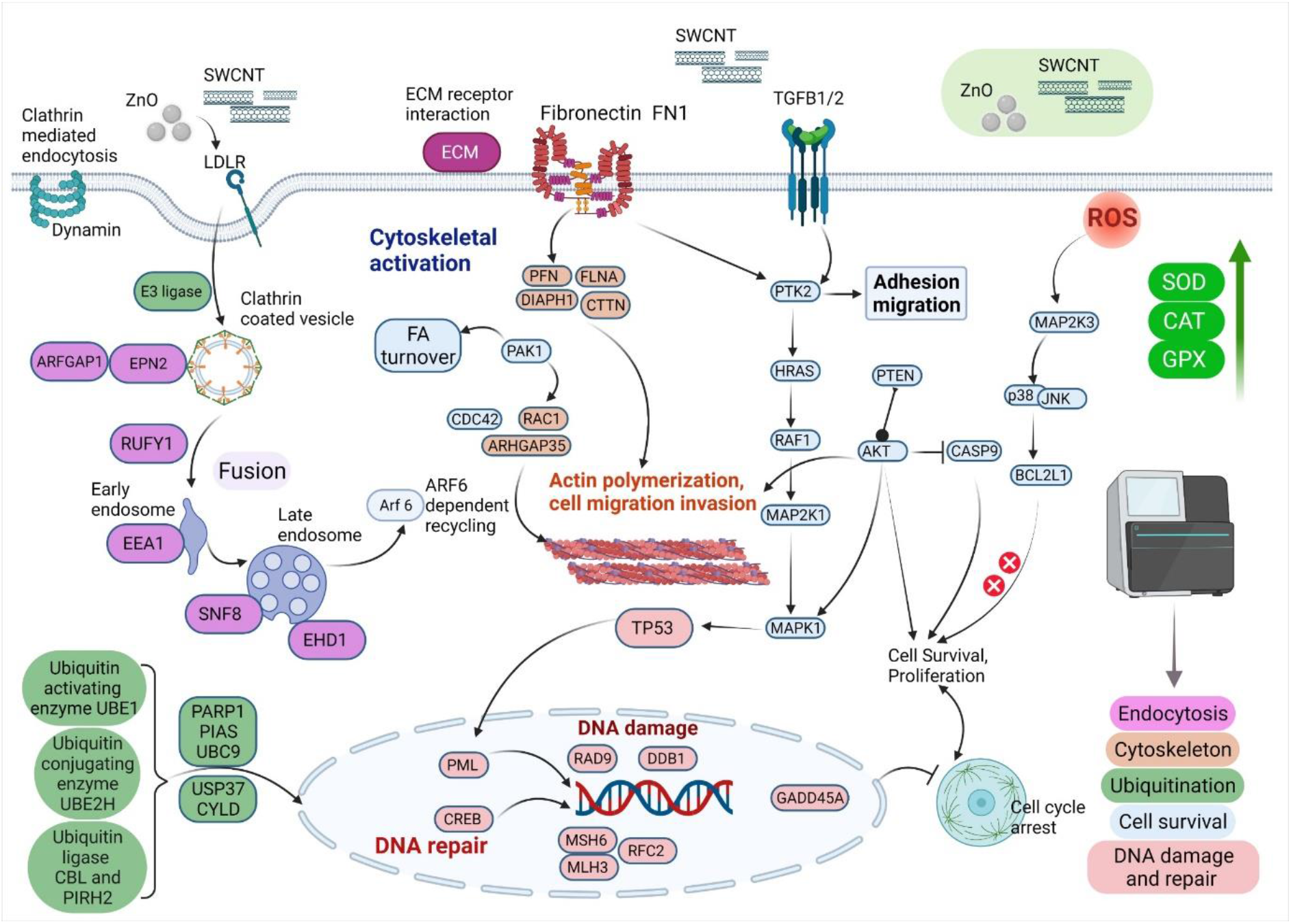
Schematic presentation of the NPs affecting the cellular process.

It was clear that in association with endocytosis, cytoskeleton remodelling, the protein ubiquitination plays a primary function in cellular homeostasis via apoptosis (Lee and Peter, 2003). Furthermore, the DNA repair mechanism and damage response connection allow the cell to counter interact with the nanoparticles.

## 4. Conclusion

In summary, we used high-throughput transcriptomics data and a liver cell line as a model system to investigate the molecular mechanism of cytotoxicity of two commonly used NPs, ZnO NPs and SWCNT. Notably, regardless of the form of NPs, low-dose (without causing major toxic effects) and acute exposure perturb related mechanistic functions. This pattern of reaction-response revealed that, regardless of NP type, the general stress response system was activated at early time points. The results showed that SWCNT and ZnO NPs reached the cells through clathrin-mediated pathways, consistent with altered cytoskeleton gene expression. The increased ROS levels within the cells may clarify the mediated DNA damage expression, but not enough to cause cell death. The results reveal the cellular response to toxic NPs and the downregulation of cell cycle-related genes. This behaviour is consistent with the findings of a cell viability assay, which showed that cell survival was sustained at >80% at low doses and early time points. Finally, we demonstrated that ZnO NPs and SWCNT activate proteasomal genes, causing ubiquitination and degradation of different proteins.

## Acknowledgement

The authors would like to thank Director ICAR-NDRI for providing sufficient funds and reagents. NCCS, Pune for giving HepG2 cell line. The authors also thank Genotypic Bengaluru for providing an RNA sequencing facility.

## Notes

### Competing Interest Statement

The authors have declared no competing interest.

